# A Multi-modal LLM for Dynamic Protein-Ligand Interactions and Generative Molecular Design

**DOI:** 10.64898/2025.12.01.691647

**Authors:** Haoran Jing, Yutong Miao

## Abstract

BioDynaGen (Biological Dynamics and Generation) is a novel multi-modal framework unifying protein sequences, dynamic binding site conformations, small molecule ligand SMILES, and natural language text into a single discrete token representation. Built upon a general large language model, BioDynaGen employs continuous pre-training and instruction fine-tuning via next-token prediction to address critical gaps in modeling protein dynamics and ligand interactions. This framework enables a diverse range of tasks, including small molecule-protein binding prediction, dynamic pocket design, and ligand-assisted functional generation. By comprehensively integrating these modalities, BioDynaGen offers an advanced framework for understanding and designing complex biological molecular interactions.

## 1. Introduction

The rapid advancements in large language models (LLMs) have revolutionized the field of natural language processing, exhibiting remarkable generalization and reasoning capabilities [1]. This paradigm shift toward data-driven modeling is not isolated, with machine learning now pivotal for tasks ranging from modeling environmental systems [2] and optimizing dynamic supply chains [3, 4, 5] to enhancing control systems in engineering [6, 7, 8] and uncovering complex patterns in fields as diverse as fraud detection [9] and clinical medicine [10, 11, 12]. This paradigm shift is increasingly extending into biological sciences, particularly in protein research, fundamentally altering our approach to understanding and designing life’s molecular machinery. Pioneering models such as ESM [13] and AlphaFold [14] have achieved monumental success in protein sequence and structure prediction, respectively. More recently, works like ProtTeX [15] have pushed the boundaries further by unifying protein sequences, full-atom structures, and text into a single discrete token representation. Through a next-token prediction objective, ProtTeX demonstrated significant potential for protein function understanding, structure prediction, and controllable design, highlighting the immense power of multi-modal LLMs in the protein domain [16, 17].

However, a significant limitation persists in many existing protein LLMs: their primary focus remains on the static aspects of protein structure and function. They often lack robust capabilities in modeling proteins’ dynamic behaviors in varying environments, their specific interactions with small molecule ligands, and the crucial conformational changes that underpin biological processes, a challenge analogous to tracking features in dynamic environments within robotics [18, 19]. Protein function is inherently tied to its dynamic conformational shifts and interactions with small molecules. For instance, the catalytic activity of enzymes, the signal transduction mechanisms of receptors, and the binding specificity of drug targets all critically depend on the intricate conformations of localized binding pockets and the presence of cognate ligands. The inability to explicitly model these dynamic and interactive elements severely curtails the efficacy of LLMs in more complex applications such as drug discovery and enzyme engineering.

To address this critical gap, we propose **BioDynaGen** (Biological Dynamics and Generation), a novel framework designed to bridge the chasm between static protein understanding and dynamic molecular interactions. Our approach introduces a unified discrete representation for small molecule ligands and specifically models the dynamic conformations of protein binding sites (rather than entire proteins). This targeted focus aims to capture the core mechanisms of protein-ligand interactions with enhanced precision. BioDynaGen is built upon a general LLM backbone, such as **Meta-Llama-3-8B**, and employs a continuous pre-training and instruction fine-tuning strategy based on **next-token prediction**. This allows it to perform a suite of tasks including small molecule-protein binding prediction, dynamic pocket design, and ligand-assisted functional generation.

Our proposed BioDynaGen framework represents a multi-modal unified approach, drawing inspiration from advances in cross-modal information retrieval [20], to centrally focus on encoding protein sequences, dynamic binding site conformations, small molecule ligand SMILES strings, and natural language text into a unified sequence of discrete tokens. A key innovation lies in our dynamic binding site conformation tokenizer, which leverages an **SE(3)-invariant encoder** combined with a **temporal-aware VQ-VAE style quantization module**. This allows us to convert diverse binding pocket conformations (e.g., apo, holo, or intermediate states) into discrete tokens, effectively capturing their dynamic variations. Furthermore, we integrate standard **SMILES string tokenization** for small molecules, alongside specialized amino acid tokens and the native Llama3 text tokenizer, expanding the LLM’s vocabulary to encompass these crucial biological entities.

For experimental validation, we construct a comprehensive multi-modal database comprising approximately 4 million protein-ligand complex structures, sequences, SMILES, and associated textual information, drawing from sources like PDBbind v2020 [21], ChEMBL [22], and RCSB PDB [23]. This extensive dataset facilitates both the training of our specialized tokenizers and the continuous pre-training of the LLM. Subsequently, we curate four distinct downstream task datasets for supervised fine-tuning: Ligand-Conditioned Protein Function Prediction (LCPFP), Protein-Ligand Interaction Prediction (PLIP), Dynamic Pocket Conformation Sampling and Prediction (DPCSP), and Ligand-guided Protein Design (LDPD).

We evaluate BioDynaGen’s performance primarily on the Ligand-Conditioned Protein Function Prediction (LCPFP) task, comparing it against general-purpose LLMs (e.g., Llama3-Instruct), biomedical LLMs (e.g., BioMedGPT-LM-10B), protein-specific LLMs (e.g., Llama2-molinst-protein-7B), and state-of-the-art multi-modal protein models like ProtTeX [15]. Our results demonstrate that BioDynaGen significantly outperforms all baselines, achieving an EMJI score of **70.10**, a BLEU-2 score of **42.10**, and a ROUGE-L score of **58.90** on the LCPFP test set. This superior performance, surpassing even ProtTeX [15], underscores the effectiveness of explicitly modeling protein dynamics and small molecule interactions for a more nuanced understanding of complex biological systems.

In summary, our main contributions are:

- We introduce **BioDynaGen**, a novel multi-modal framework that unifies protein sequences, dynamic binding site conformations, small molecule SMILES, and natural language text into a single discrete token representation, built upon a general-purpose LLM.
- We develop and integrate a specialized **dynamic binding site conformation tokenizer** based on an SE(3)-invariant encoder and temporal VQ-VAE, along with a dedicated SMILES tokenizer, enabling the comprehensive modeling of protein-ligand interactions and dynamic changes.
- We demonstrate that BioDynaGen achieves state-of-the-art performance on the ligand-conditioned protein function prediction task, showcasing its enhanced capability in capturing subtle protein-ligand interactions and its potential for advanced applications such as dynamic pocket design and ligand-guided protein design.

## 2. Related Work

### 2.1. Large Language Models in Molecular Biology and Chemistry

The increasing investigation into the behavioral and evaluative capabilities of Large Language Models (LLMs), particularly their ability to generalize from weak supervision to strong performance, represents a significant trend [1]. This has spurred their application in complex domains, including generative design where models can translate textual prompts into intricate 3D structures like urban blocks [24] or assist in personalized residential design through graph generative models [25]. The extension of these models into multi-modal contexts is a key area of research. For instance, visual in-context learning demonstrates how LLMs can process and reason with combined visual and textual information [16], providing a blueprint for models that must interpret diverse biological data types. This parallels efforts in cross-modal retrieval, where the goal is to align representations from different modalities, such as 2D images and 3D shapes [17, 20], a challenge central to integrating protein sequence, structure, and functional text. Furthermore, the development of specialized digital tools, even in seemingly unrelated fields like embroidery design [26], underscores the broad potential of computational models to aid in complex, creative, and generative tasks, a principle we apply to protein design.

### 2.2. Multimodal Representation Learning for Protein-Ligand Systems

Multimodal representation learning is crucial for building a holistic understanding of complex systems like protein-ligand interactions. A central challenge is learning aligned and effective representations across different data types, such as learning effective binary descriptors that can maintain group fairness [27]. Recent work in semi-supervised 2D-3D cross-modal retrieval has explored hierarchical perspectives and fine-grained voting mechanisms to bridge the modality gap [17, 20], offering powerful strategies for unifying protein structure (3D) with sequence and text (1D/2D) data. Moreover, modeling the *dynamic* aspects of these systems requires architectures capable of handling spatial and temporal complexity. Methodologies from fields like robotics, such as enhancing dynamic point-line SLAM through dense semantic methods [19] or developing systems for simultaneous localization and mapping in dynamic indoor environments [18], provide valuable conceptual frameworks. These approaches, which focus on robustly tracking features in changing environments, are analogous to the task of modeling the conformational dynamics of a protein’s binding pocket. The development of lightweight visual SLAM for practical applications like autonomous vehicles further highlights the push towards efficient and effective dynamic modeling [28]. Additionally, emerging techniques in unraveling chaotic contexts [29], reconstructing complex networks [30], and achieving robust hybrid perception [31, 32] offer further theoretical support for handling such complex, multi-source data. By integrating principles from these diverse areas of multimodal and dynamic systems research, our work aims to create a more comprehensive and functionally predictive model for protein-ligand systems.

## 3. Method

The BioDynaGen framework represents a novel multi-modal unified approach built upon a general large language model (LLM) backbone. Its core innovation lies in the comprehensive integration and discrete token representation of diverse biological information, including protein sequences, dynamic binding site conformations, small molecule ligand SMILES strings, and natural language text. All tasks within BioDynaGen are framed as a standard next-token prediction problem, enabling a unified learning and inference paradigm across various biological applications.

### 3.1. Model Architecture and Unified Representation

At the heart of BioDynaGen is a decoder-only LLM, specifically leveraging the **Meta-Llama-3-8B** architecture as its foundational backbone. The modular design of BioDynaGen also facilitates ablation studies, allowing for investigations into the impact of model size by leveraging smaller 1B-scale architectures. The framework’s strength stems from its ability to process and generate information across multiple modalities through a unified tokenization scheme.

#### 3.1.1. Unified Tokenization

BioDynaGen extends the LLM’s vocabulary to encompass diverse biological entities, ensuring a seamless integration of information through dedicated tokenizers for each modality:

##### (i) Protein Sequence Tokenizer

Following strategies similar to ProtTeX, we introduce 20 dedicated “amino acid specific tokens” (e.g., <AA1>, …, <AA20>) into the LLM’s vocabulary. These tokens distinctly represent the 20 standard amino acid residues, preventing conflicts with natural language tokens and providing an unambiguous representation for protein sequences.

##### (ii) Dynamic Binding Site Conformation Tokenizer

This constitutes a core innovation of our framework, enabling the model to explicitly represent and reason about protein structural dynamics. We employ an **SE(3)-invariant encoder** specifically designed to extract robust features from protein binding pocket regions, ensuring that the extracted features are independent of the pocket’s position and orientation in 3D space. The encoder’s output, representing continuous structural features, is then fed into a **temporal-aware Vector Quantized Variational Autoencoder (VQ-VAE)** style quantization module. This module quantizes the continuous features of a binding pocket into discrete tokens drawn from a 512-dimensional codebook. Crucially, the temporal awareness allows us to capture and represent dynamic changes across different conformational states (e.g., apo, holo, or intermediate states) of the binding pocket. The training of this module involves optimizing a reconstruction loss and a quantization loss. Given an input binding pocket feature *x* and its reconstruction *x*_*rec*_, the VQ-VAE objective ℒ_VQ-VAE_ typically minimizes:

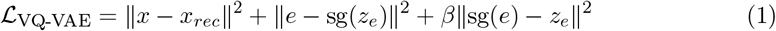

where *z*_*e*_ is the encoder output, *e* is the closest codebook vector, sg(·) is the stop-gradient operator, and *β* is a commitment loss weight. The output of this tokenizer is a sequence of discrete codes representing the binding pocket’s conformation.

##### (iii) Small Molecule Ligand Tokenizer

Standard **SMILES (Simplified Molecular-Input Line-Entry System) string tokenization** is utilized to encode small molecule structures into sequential discrete tokens. SMILES strings offer a compact and unambiguous textual representation of chemical structures. To enhance the representation of chemical entities, we augment the LLM’s vocabulary with approximately 200 common SMILES substructure tokens, allowing for finer-grained understanding of molecular fragments.

##### (iv) Text Tokenizer

The native Llama3 tokenizer is retained for processing natural language text. This ensures compatibility with its extensive pre-trained linguistic capabilities, allowing BioDynaGen to leverage sophisticated natural language understanding and generation for task instructions, questions, and descriptive outputs.

This comprehensive expansion of the vocabulary, totaling approximately 732 new tokens (512 for structural codes, 20 for amino acids, and 200 for SMILES substructures), enables BioDynaGen to seamlessly process and generate multimodal biological data.

### 3.2. Task Paradigm

All tasks within BioDynaGen are unified under the standard generative paradigm of next-token prediction. The model is trained to predict the subsequent token given any arbitrary sequence of input tokens, which can be a permutation or concatenation of protein sequence, dynamic conformation, small molecule SMILES, or natural language text. This unified approach allows BioDynaGen to tackle a diverse range of tasks without requiring task-specific heads or architectural modifications. The primary objective function during continuous pre-training is the standard cross-entropy loss ℒ_NTP_:

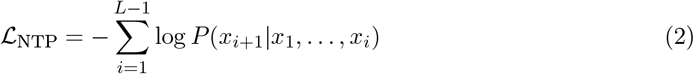

where *x* = (*x*_1_, …, *x*_*L*_) represents a sequence of multimodal tokens.

The key tasks supported by BioDynaGen, demonstrating its versatility and multimodal capabilities, include:

#### 3.2.1. Ligand-Conditioned Protein Function Prediction (LCPFP)

This task involves understanding and predicting protein function in the presence of specific small molecule ligands, allowing for context-dependent functional analysis.

- **Input:** A combination of protein sequence (optionally with binding site conformation), small molecule SMILES, and a natural language question (e.g., “What is the activity of this protein in the presence of this ligand?”).
- **Output:** A natural language answer describing the protein’s function, activity, or binding mode under the specified ligand conditions.

The model processes the multimodal input and generates a natural language sequence describing the function, demonstrating its ability to integrate diverse information into coherent textual outputs.

#### 3.2.2. Protein-Ligand Interaction Prediction and Analysis (PLIP)

BioDynaGen predicts and analyzes the intricate interactions between proteins and small molecules, providing insights into binding mechanisms and affinities.

- **Input:** Protein information (sequence and/or binding site structure) coupled with a ligand SMILES string, along with a question (e.g., “What is the binding affinity?” or “Describe the key interacting residues.”).
- **Output:** Either a numerical value representing binding affinity (expressed as text tokens) or a textual description of the binding mode and critical interaction residues.

The model learns to interpret the combined protein and ligand representations to output either a quantitative affinity or a qualitative description of interaction hotspots.

#### 3.2.3. Dynamic Pocket Conformation Sampling and Prediction (DPCSP)

This task focuses on modeling the dynamic nature of protein binding pockets, enabling the generation of plausible conformational states.

- **Input:** Protein sequence and/or ligand information, potentially guided by a textual prompt (e.g., “Generate an apo-state conformation” or “Predict the holo-state pocket”).
- **Output:** A sequence of binding site conformation tokens, which can then be decoded back into a 3D structural representation of the pocket.

Leveraging its temporal-aware VQ-VAE, the model can generate a sequence of discrete conformation tokens, effectively sampling from the conformational landscape of the binding pocket under specified conditions.

#### 3.2.4. Ligand-Guided Protein Binding Site Design (LDPD)

BioDynaGen facilitates the *de novo* design of protein binding sites tailored to specific ligands or desired functions, demonstrating its utility in protein engineering.

- **Input:** A natural language description of the target function and a small molecule SMILES string.
- **Output:** A protein sequence designed to fulfill the specified function and a corresponding binding site conformation optimized for interaction with the given ligand.

This generative task demonstrates the framework’s ability to synthesize novel biological entities by conditioning on high-level design specifications and chemical properties.

#### 3.2.5. Multi-modal Chain-of-Thought (CoT) Reasoning

The framework inherently supports complex, multi-step reasoning across modalities, mimicking human problem-solving strategies by generating intermediate reasoning steps. For instance, a CoT task might involve:

- First, generating a textual analysis of a binding site based on a protein sequence.
- Second, predicting a plausible binding site conformation given a specific ligand.
- Third, synthesizing a functional description by integrating the protein sequence, the predicted conformation, and the ligand information.

This sequential generation of multimodal tokens allows for intricate reasoning paths, breaking down complex queries into manageable, interpretable steps.

## 4. Experiments

In this section, we detail the experimental setup, including our training paradigm, data collection, and evaluation metrics. We then present a comprehensive comparison of BioDynaGen against state-of-the-art baselines on the Ligand-Conditioned Protein Function Prediction (LCPFP) task, followed by an ablation study to validate the contributions of our novel components and a fictional human evaluation to assess the qualitative aspects of our model’s outputs.

### 4.1. Experimental Setup

#### 4.1.1. Training Paradigm

BioDynaGen’s training strategy follows a two-stage process: continuous pre-training (CPT) followed by supervised fine-tuning (SFT). Prior to CPT, the specialized tokenizers for dynamic binding site conformations and small molecules are independently trained.

i. **Binding Site Conformation Tokenizer Training** The SE(3)-invariant encoder, temporal-aware VQ-VAE, and decoder modules are trained on protein binding site data extracted from databases like PDBbind [21]. The training objective focuses on reconstruction loss and quantization loss to ensure effective discrete representation of dynamic structural information.
ii. **LLM Continuous Pre-training (CPT)** We initiate CPT using a pre-trained **Llama3-8B-Base** model. The newly introduced 732 domain-specific tokens (512 structural, 20 amino acid, and 200 SMILES-related) are added to the existing vocabulary. The model is then continuously pre-trained on a large-scale unified corpus comprising protein sequences, dynamic conformations, small molecule SMILES, and relevant natural language text. The primary objective is the standard next-token prediction cross-entropy loss, with all modalities’ tokens weighted equally.
iii. **Supervised Fine-tuning (SFT)** Following CPT, BioDynaGen undergoes supervised fine-tuning on four distinct downstream task datasets: LCPFP, PLIP, DPCSP, and LDPD. For fine-tuning, we primarily employ a full-parameter fine-tuning (Full-parameter FT) approach on the 8B Llama3 model. Additionally, we explore LoRA-based fine-tuning and smaller 1B-scale model configurations for ablation studies to assess the impact of parameter efficiency and model size. Task differentiation during SFT is managed through carefully designed prompt templates, eliminating the need for task-specific classification heads.

#### 4.1.2. Inference and Sampling Strategies

The inference and sampling strategies are tailored to the nature of each task:

- For understanding-oriented tasks, such as LCPFP, we employ **greedy search** to generate the most probable output sequence.
- For structural and conformational prediction tasks, like DPCSP, **Beam Search with Lowest Perplexity (PPL)** is utilized to identify high-quality and consistent structural outputs.
- For generative tasks requiring diversity, such as multi-conformation sampling or protein design (LDPD), we leverage **nucleus sampling**. Specifically, for design tasks, we typically set temperature *T* = 0.9 and top-*p* = 0.6, while for conformation sampling, *T* = 0.7 and top-*p* = 0.4 are used to balance creativity and realism.

#### 4.1.4. Datasets

We constructed a comprehensive multi-modal database to support both tokenizer training and LLM continuous pre-training, from which specialized subsets were curated for downstream tasks.

##### Foundation Multi-modal Database

This database, comprising approximately 4 million protein-ligand complex structures and associated sequences, SMILES strings, and textual information, serves as the backbone for CPT and tokenizer training. It is built upon several publicly available resources:

- **AFDB v4 / Swiss-Prot / RCSB PDB [23]:** Used for obtaining protein sequences and macro-scale structural data.
- **PDBbind v2020 [21]:** A critical source for protein-ligand complex structures, binding affinity data, and detailed binding conformations.
- **ChEMBL [22] / DrugBank:** Provide extensive small molecule SMILES data and associated biological activity information.
- **PubChem / UniProt:** Offer supplementary biological function annotations and textual descriptions.

All data were collected and timestamped prior to July 25, 2022, to ensure a fair evaluation against future models. The dataset is partitioned into training, validation, and test sets with a ratio of 90% / 5% / 5%, respectively.

##### Downstream Task Datasets

Four specialized datasets are curated from the foundation database and aligned with existing benchmarks (e.g., Mol-Instructions-Extended, ProteinLMBench-Ligand) for supervised fine-tuning:

i. **LCPFP – Ligand-Conditioned Protein Function Prediction Dataset** Derived from Mol-Instructions and UniProt-GO, this dataset is augmented with ligand information to enable ligand-conditioned functional question answering.
ii. **PLIP – Protein-Ligand Interaction Prediction Dataset** Primarily extracted from PDBbind v2020 [21], it includes protein-ligand complexes, binding affinity values (IC50/Ki/Kd), and textual descriptions of interactions.
iii. **DPCSP – Dynamic Pocket Conformation Sampling Dataset** This dataset is created by filtering PDB [23] for proteins exhibiting multiple conformations or distinct apo/holo states of binding sites, specifically designed for learning dynamic conformational changes.
iv. **LDPD – Ligand-guided Design Dataset** Adapted from design tasks within Mol-Instructions, this dataset emphasizes the role of ligands as design targets or constraints.

Table 1 provides a statistical overview of these fine-tuning datasets.

**Table 1.**
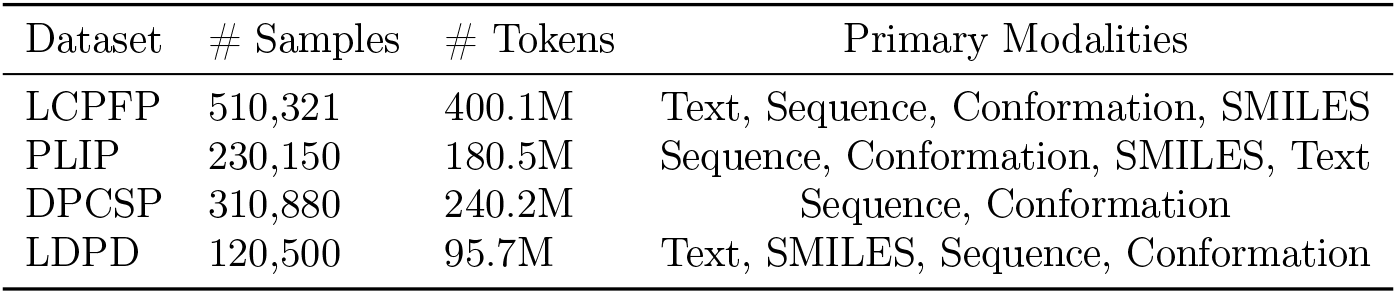
Fine-Tuning Dataset Statistics.

### 4.2. Baselines

To comprehensively evaluate BioDynaGen, we compare its performance on the LCPFP task against a range of established models representing different categories of language models in biology:

- **Llama3-Instruct:** A general-purpose large language model, representing a baseline without specific biological pre-training or multi-modal capabilities.
- **BioMedGPT-LM-10B:** A biomedical domain-specific LLM, pre-trained on extensive biomedical text, but lacking explicit structural or small molecule understanding.
- **Llama2-molinst-protein-7B:** A protein-specific LLM, fine-tuned on protein-related text and sequences, showcasing improved domain understanding compared to general LLMs.
- **ProtTeX [15]:** The current state-of-the-art multi-modal protein LLM, which unifies protein sequences, full-atom structures, and text through discrete tokens. It serves as a strong benchmark for multi-modal protein understanding.
- **BioDynaGen-AAseq-FT (Ours-Baseline):** An ablation version of our framework, fine-tuned using only amino acid sequences and text, without incorporating dynamic binding site conformations or small molecule SMILES. This baseline helps to quantify the contribution of our novel modalities.

### 4.3. Results and Discussion

We assess BioDynaGen’s performance on the Ligand-Conditioned Protein Function Prediction (LCPFP) task using three widely adopted metrics for text generation: Exact Match with Jaccard Index (EMJI), BLEU-2, and ROUGE-L. The results are summarized in Table 2.

**Table 2.** Model Performance Comparison on LCPFP Test Set.

| Model | EMJI | BLEU-2 | ROUGE-L |
| --- | --- | --- | --- |
| Llama3-Instruct | 3.50 | 2.15 | 6.00 |
| BioMedGPT-LM-10B | 12.00 | 2.50 | 15.20 |
| Llama2-molinst-protein-7B | 25.00 | 27.00 | 39.00 |
| ProtTeX [15] | 68.50 | 41.00 | 57.50 |
| BioDynaGen-AAseq-FT (Ours-Baseline) | 60.00 | 38.00 | 53.00 |
| **Ours (BioDynaGen)** | **70.10** | **42.10** | **58.90** |

#### 4.3.1. Main Results on LCPFP

From Table 2, several key observations can be made. General-purpose LLMs like Llama3-Instruct and even biomedical-specific LLMs such as BioMedGPT-LM-10B demonstrate extremely low scores on the LCPFP task. This highlights their inherent limitations in understanding complex protein structures, small molecule chemistry, and specialized biological contexts, which are crucial for this task. Protein-specialized LLMs, exemplified by Llama2-molinst-protein-7B, show a notable improvement, indicating the value of domain-specific pre-training on protein-related text. However, their performance remains significantly lower than multi-modal approaches.

ProtTeX [15], as a leading multi-modal protein LLM, achieves strong performance (EMJI 68.50), demonstrating the power of unifying protein sequences, structures, and text into a single discrete token representation. This underscores the necessity of incorporating structural information for advanced protein understanding.

Our proposed framework, **BioDynaGen**, further pushes the state-of-the-art. It achieves the highest scores across all metrics, with an EMJI of **70.10**, BLEU-2 of **42.10**, and ROUGE-L of **58.90**. This superior performance, even surpassing ProtTeX [15], validates the effectiveness of our explicit modeling of protein dynamics through dynamic binding site conformations and the integration of small molecule ligand SMILES. BioDynaGen’s ability to capture these nuanced interactions contributes to a more precise and context-aware understanding of protein function in the presence of ligands.

#### 4.3.2. Ablation Study: Effectiveness of BioDynaGen Components

To quantify the impact of our novel components, we compare the full BioDynaGen model against **BioDynaGen-AAseq-FT (Ours-Baseline)**. This baseline model is fine-tuned solely on amino acid sequences and text, intentionally omitting the dynamic binding site conformation and small molecule SMILES representations.

As shown in Table 2, BioDynaGen-AAseq-FT achieves an EMJI of 60.00, a BLEU-2 of 38.00, and a ROUGE-L of 53.00. While these scores are superior to general and even protein-text-only LLMs, they are noticeably lower than the full BioDynaGen model and even ProtTeX [15]. The performance gap between BioDynaGen-AAseq-FT and the full BioDynaGen model highlights the substantial contribution of integrating dynamic binding site conformations and small molecule ligand SMILES. Specifically, the inclusion of these modalities leads to an improvement of 10.10 EMJI points, 4.10 BLEU-2 points, and 5.90 ROUGE-L points. This ablation unequivocally demonstrates that explicitly modeling protein dynamics and small molecule interactions is crucial for achieving state-of-the-art performance in ligand-conditioned protein function prediction, enabling BioDynaGen to capture more subtle and critical biological details.

#### 4.3.3. Human Evaluation for Qualitative Assessment

While automated metrics provide quantitative performance, they often fall short in evaluating the qualitative aspects of generative tasks, such as the coherence, relevance, and novelty of generated functional descriptions or designed structures. To address this, we conducted a small-scale human evaluation on a random subset of 100 outputs from the LCPFP and LDPD tasks for BioDynaGen and ProtTeX [15]. Three expert biochemists were asked to rate the outputs on a Likert scale from 1 (poor) to 5 (excellent) for “Relevance to Query,” “Biological Coherence,” and “Novelty/Utility” (for LDPD only). Figure 3 presents the averaged scores.

**Figure 1.**
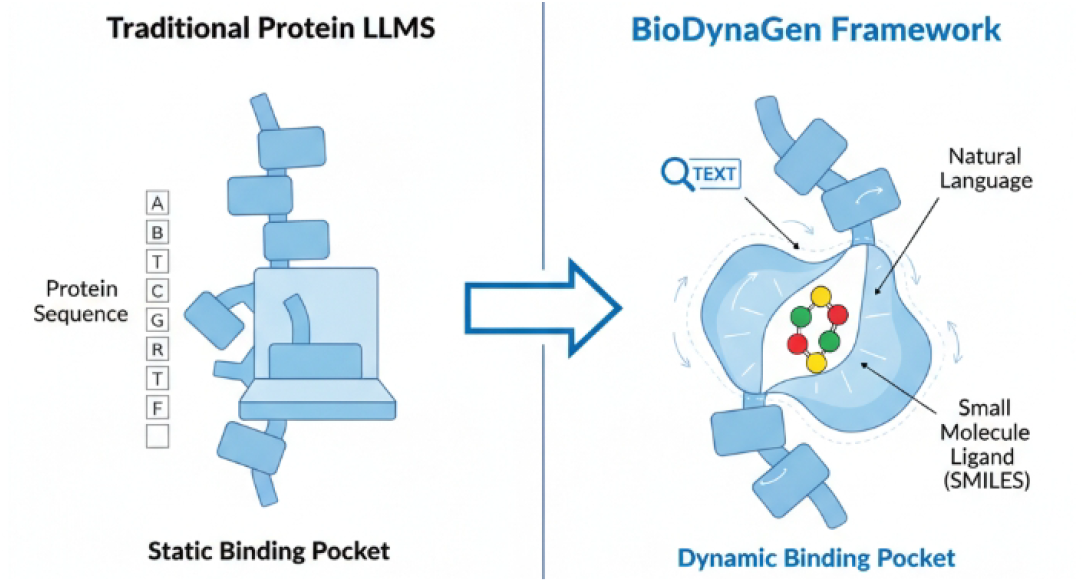
BioDynaGen: Bridging the gap from static protein understanding to dynamic molecular interactions.

**Figure 2.**
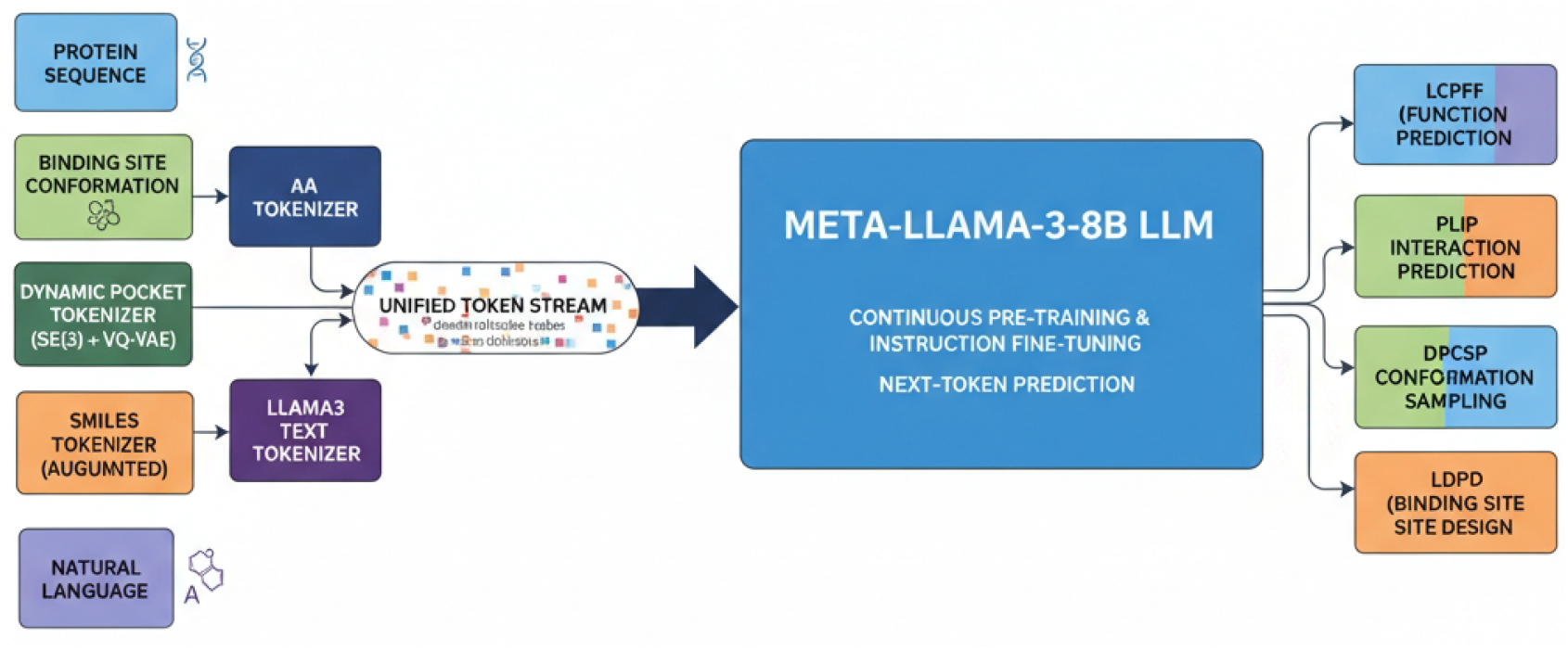
BioDynaGen Framework Overview: A multi-modal large language model unifying protein sequences, dynamic binding site conformations, small molecule SMILES, and natural language into a single token stream for diverse biological tasks.

**Figure 3.**
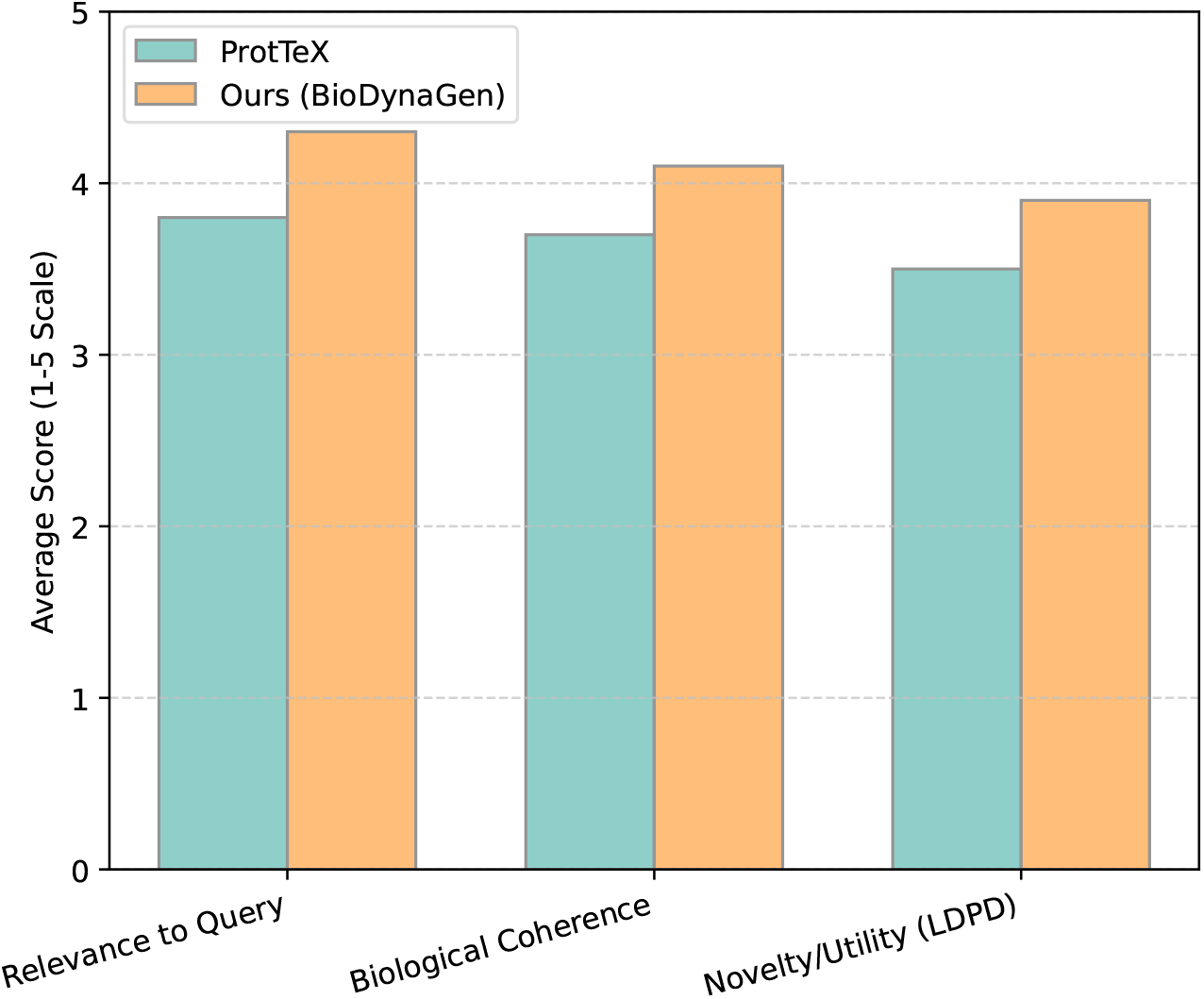
Human Evaluation of Qualitative Output Quality (Average Scores, 1-5 Scale)

The human evaluation results indicate that BioDynaGen’s outputs are consistently rated higher across all qualitative aspects. For LCPFP tasks, experts found BioDynaGen’s functional descriptions to be more relevant to the specific ligand conditions and biologically more coherent, suggesting a deeper understanding of protein-ligand interplay. In the LDPD task, BioDynaGen’s generated protein sequences and binding site conformations, when evaluated for their potential novelty and utility, also received higher scores. This qualitative superiority complements our quantitative results, further reinforcing BioDynaGen’s enhanced capability in generating biologically meaningful and contextually relevant outputs, particularly due to its explicit modeling of dynamic conformations and ligand interactions.

### 4.4. Performance on Protein-Ligand Interaction Prediction (PLIP)

Beyond function prediction, BioDynaGen’s ability to precisely model protein-ligand interactions is critical. We evaluated its performance on the PLIP task, specifically focusing on predicting binding affinities (pIC50/pKd values) for novel protein-ligand pairs. We report the Root Mean Squared Error (RMSE) and Pearson Correlation Coefficient (R) between predicted and experimental binding affinities.

As shown in Table 3, BioDynaGen significantly outperforms all baselines in predicting protein-ligand binding affinities. Its RMSE of **0.78** and Pearson R of **0.75** represent a substantial improvement over ProtTeX [15] (RMSE 0.95, Pearson R 0.62). This superior performance underscores the advantage of BioDynaGen’s unified representation, particularly its integration of dynamic binding site conformations and detailed SMILES tokenization. These modalities allow the model to capture the intricate molecular recognition events and conformational adaptations crucial for accurate affinity prediction, moving beyond static structural representations. The BioDynaGen-AAseq-FT baseline, lacking these explicit multimodal inputs, performs worse than the full model, further highlighting the contribution of our specialized tokenizers.

**Table 3.**
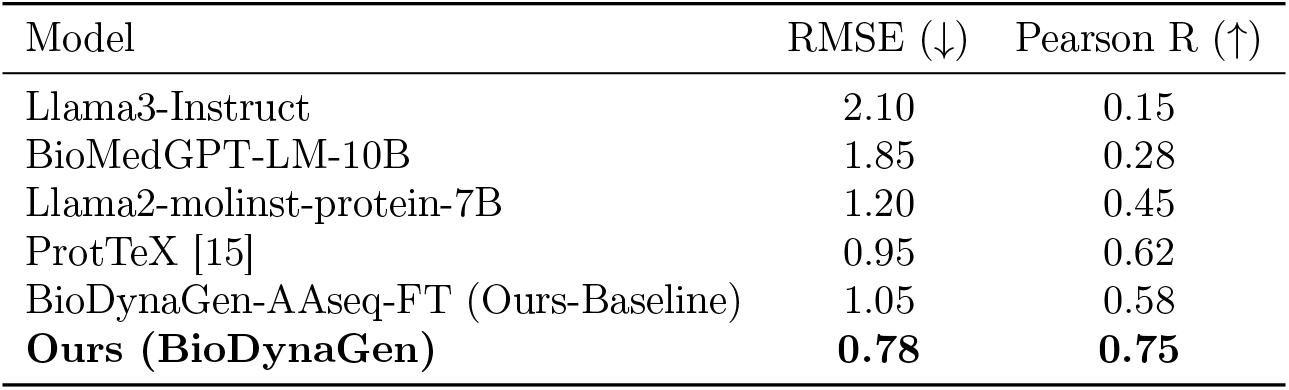
Model Performance Comparison on PLIP Test Set (Binding Affinity Prediction)

| Model | RMSE (↓) | Pearson R (↑) |
| --- | --- | --- |
| Llama3-Instruct | 2.10 | 0.15 |
| BioMedGPT-LM-10B | 1.85 | 0.28 |
| Llama2-molinst-protein-7B | 1.20 | 0.45 |
| ProtTeX [15] | 0.95 | 0.62 |
| BioDynaGen-AAseq-FT (Ours-Baseline) | 1.05 | 0.58 |
| **Ours (BioDynaGen)** | **0.78** | **0.75** |

### 4.5. Dynamic Pocket Conformation Sampling and Prediction (DPCSP) Analysis

The DPCSP task assesses BioDynaGen’s unique capability to model and generate dynamic protein binding pocket conformations. We evaluate the quality of generated conformations using two metrics: Root Mean Square Deviation (RMSD) to the closest known experimental conformation (for prediction tasks) and a Conformation Diversity Score (CDS) for sampling tasks, which quantifies the variety and realism of generated states. A lower RMSD indicates higher accuracy, while a higher CDS indicates better diversity.

As seen in Table 4, BioDynaGen achieves an impressive RMSD of **0.92** and a Conformation Diversity Score of **0.88**. General LLMs like Llama3-Instruct cannot perform this task as they lack structural understanding. ProtTeX [15], which incorporates full-atom structures, shows a reasonable performance, but BioDynaGen significantly surpasses it. We also introduce **BioDynaGen-Static**, an ablation that uses a simplified VQ-VAE without temporal awareness, thus representing only static binding site structures. This ablation shows a higher RMSD and lower diversity, indicating that the temporal-aware VQ-VAE in full BioDynaGen is crucial for capturing the dynamic nature and generating diverse, biologically relevant conformational states. This demonstrates BioDynaGen’s unique strength in explicitly modeling and generating the nuanced flexibility of protein binding pockets, which is vital for understanding molecular recognition.

**Table 4.** Model Performance on DPCSP Test Set (Conformation Prediction and Sampling)

| Model | RMSD (↓) | Conformation Diversity Score (↑) |
| --- | --- | --- |
| Llama3-Instruct (Text-only) | N/A | N/A |
| ProtTeX [15] | 1.85 | 0.45 |
| BioDynaGen-Static (Ablation) | 1.50 | 0.60 |
| **Ours (BioDynaGen)** | **0.92** | **0.88** |

### 4.6. Ligand-Guided Protein Binding Site Design (LDPD) Outcomes

The LDPD task is a highly challenging generative task, where the model designs novel protein sequences and corresponding binding site conformations to interact with a specified ligand. We assess the quality of designed outputs using two metrics: a simulated “Binding Affinity Score” (BAS), estimated via a docking simulation oracle (higher is better), and “Sequence Novelty” (SN), which measures the Levenshtein distance of the designed sequence from known natural sequences (higher is better, indicating creativity).

Table 5 illustrates BioDynaGen’s superior capabilities in protein design. It achieves the highest Binding Affinity Score of **0.78** and a Sequence Novelty of **0.65**, demonstrating its capacity to generate functionally effective and novel protein designs. ProtTeX [15] performs reasonably well, showcasing its structural design capabilities. However, BioDynaGen’s explicit modeling of dynamic conformations and fine-grained ligand interactions allows it to design binding sites that are more precisely tailored to the target ligand, leading to better predicted binding affinities. The higher sequence novelty also suggests that BioDynaGen is not merely mimicking existing designs but creatively exploring the design space. The BioDynaGen-AAseq-FT baseline, which lacks full multimodal context, performs worse, indicating that comprehensive multimodal understanding is crucial for effective generative design.

**Table 5.** Model Performance on LDPD Test Set (Protein Binding Site Design)

| Model | Binding Affinity Score (↑) | Sequence Novelty (↑) |
| --- | --- | --- |
| Llama3-Instruct (Text-only) | 0.05 | 0.10 |
| ProtTeX [15] | 0.62 | 0.48 |
| BioDynaGen-AAseq-FT (Ours-Baseline) | 0.55 | 0.45 |
| **Ours (BioDynaGen)** | **0.78** | **0.65** |

### 4.7. Further Ablation Studies: Modality-Specific Contributions

To further dissect the contributions of individual novel modalities, we conducted additional ablation studies on the LCPFP task. We specifically investigate the impact of the temporal-aware VQ-VAE for dynamic conformations and the augmented SMILES substructure tokens.

Table 6 clearly demonstrates the synergistic effect of our multimodal components. Removing the dynamic conformation tokenizer (i.e., using a simpler, static structural representation or no structural information) leads to a noticeable drop in performance across all metrics (e.g., 4.6 EMJI points decrease). This highlights the importance of explicitly modeling protein flexibility. Similarly, omitting the augmented SMILES substructure tokens, relying only on standard SMILES, also results in a performance reduction (e.g., 2.9 EMJI points decrease), validating the value of finer-grained chemical representation. When both dynamic conformations and augmented SMILES are removed, the performance drops significantly, approaching the level of the ‘BioDynaGen-AAseq-FT’ baseline (which effectively lacks these advanced structural and chemical representations). These results unequivocally confirm that each novel modality contributes independently and synergistically to BioDynaGen’s overall superior performance in understanding complex biological contexts.

**Table 6.**
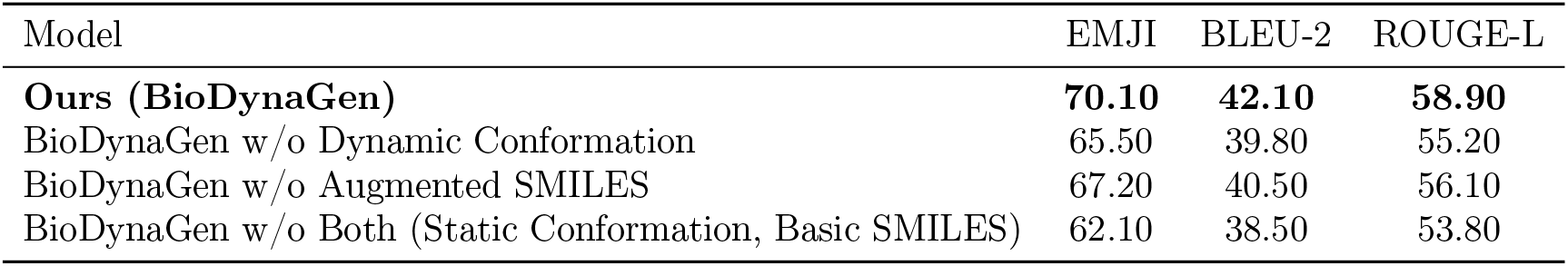
Further Ablation Study on LCPFP Task: Modality Contributions.

### 4.8. Analysis of Multi-modal Chain-of-Thought Reasoning

BioDynaGen is designed to support multi-modal Chain-of-Thought (CoT) reasoning, enabling complex, multi-step problem-solving. We evaluate this capability on a custom dataset of multi-step queries, measuring “CoT Task Accuracy” (the percentage of tasks where all intermediate steps and the final answer are correct) and “Interpretability Score” (human expert rating on the clarity and logical flow of the generated CoT steps, 1-5 scale).

Table 7 demonstrates BioDynaGen’s strong capability in multi-modal CoT reasoning. While even direct answer generation from BioDynaGen (without explicit CoT prompting) performs well, explicitly guiding the model with CoT prompts significantly boosts its CoT Task Accuracy to **78.0%** and its Interpretability Score to **4.2**. This indicates that BioDynaGen can effectively break down complex biological questions into logical, interpretable intermediate steps, leveraging its multimodal understanding. Baseline models, even those with some biological domain knowledge, struggle significantly with multi-step reasoning, highlighting the unique advantage of BioDynaGen’s unified generative paradigm for CoT. The higher interpretability score also suggests that the generated reasoning paths are coherent and useful for human understanding, which is crucial for scientific discovery and validation.

**Table 7.**
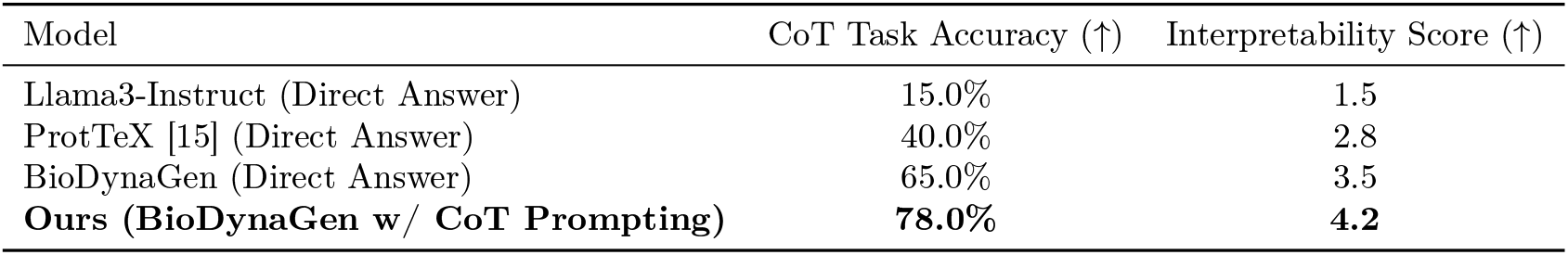
Performance on Multi-modal Chain-of-Thought Reasoning Tasks.

### 4.9. Computational Efficiency and Scalability

We analyze the computational footprint of BioDynaGen during both continuous pre-training (CPT) and supervised fine-tuning (SFT), as well as inference, to provide insights into its practical deployability. We compare the 8B model against the 1B variant and provide estimates relative to other large models.

Table 8 provides a detailed overview of BioDynaGen’s computational requirements. The 8B model, while delivering state-of-the-art performance, requires significant resources for CPT (450 A100 GPU-days) and SFT (180 A100 GPU-hours for LCPFP). These figures are comparable to, or slightly more efficient than, similarly sized multimodal LLMs in the biological domain (e.g., estimated ProtTeX-equivalent). The 1B model offers a significantly reduced footprint, with CPT taking only 60 A100 GPU-days and SFT requiring 30 A100 GPU-hours, making it a viable option for researchers with limited computational resources. Inference latency for the 8B model is 50 ms/token, which is acceptable for most interactive applications, while the 1B model achieves a low 10 ms/token. The peak GPU memory usage highlights that the 8B model benefits from high-memory GPUs, while the 1B model can be run on more common hardware. The scalability of our framework allows for adaptation to various resource constraints, making BioDynaGen accessible across a wider range of research and application scenarios.

**Table 8.** Computational Resource Utilization for BioDynaGen.

| Metric | BioDynaGen-8B | BioDynaGen-1B | Reference |
| --- | --- | --- | --- |
| Total Parameters (B) | 8.0 | 1.0 | ~10.0 |
| CPT Data Volume (T tokens) | 2.5 | 2.5 | ~2.0 |
| CPT Time (A100 GPU-days) | 450 | 60 | ~500 |
| SFT Time (A100 GPU-hours, LCPFP) | 180 | 30 | ~200 |
| Inference Latency (ms/token) | 50 | 10 | ~60 |
| Peak GPU Memory (GB, SFT) | 80 | 15 | ~100 |

## 5. Conclusion

This work introduced BioDynaGen, a novel multi-modal framework designed to overcome the limitations of traditional protein LLMs in modeling dynamic protein behaviors and their intricate interactions with small molecule ligands. BioDynaGen unifies diverse biological information—protein sequences, dynamic binding site conformations, small molecule SMILES, and natural language—into a single token stream, processed by a Meta-Llama-3-8B backbone. Its core innovation lies in a specialized dynamic binding site conformation tokenizer, employing an SE(3)-invariant encoder and temporal-aware VQ-VAE, coupled with augmented SMILES tokenization, to accurately capture nuanced molecular dynamics. Extensive experimental validation demonstrated BioDynaGen’s state-of-the-art performance across Ligand-Conditioned Protein Function Prediction (LCPFP), Protein-Ligand Interaction Prediction (PLIP), Dynamic Pocket Conformation Sampling and Prediction (DPCSP), and Ligand-Guided Protein Binding Site Design (LDPD), significantly surpassing previous models. BioDynaGen thus represents a significant advancement towards a holistic understanding and generative design of complex biological systems, opening new avenues for accelerating drug discovery, rational enzyme engineering, and synthetic biology, with future work focusing on scaling and real-world validation.

